# Differential integration of transcriptome and proteome identifies pan-cancer prognostic biomarkers

**DOI:** 10.1101/322313

**Authors:** Gregory W. Schwartz, Jelena Petrovic, Yeqiao Zhou, Robert B. Faryabi

## Abstract

High-throughput analysis of the transcriptome and proteome individually are used to interrogate complex oncogenic processes in cancer. However, an outstanding challenge is how to combine these complementary, yet partially disparate data sources to accurately identify tumor-specific gene-programs and clinical biomarkers. Here, we introduce `inteGREAT` for robust and scalable differential integration of high-throughput measurements. With `inteGREAT`, each data source is represented as a co-expression network, which is analyzed to characterize the local and global structure of each node across networks. `inteGREAT` scores the degree by which the topology of each gene in both transcriptome and proteome networks are conserved within a tumor type, yet different from other normal or malignant cells. We demonstrated the high performance of `inteGREAT` based on several analyses: deconvolving synthetic networks, rediscovering known diagnostic biomarkers, establishing relationships between tumor lineages, and elucidating putative prognostic biomarkers which we experimentally validated. Furthermore, we introduce the application of a clumpiness measure to quantitatively describe tumor lineage similarity. Together, `inteGREAT` not only infers functional and clinical insights from the integration of transcriptomic and proteomic data sources in cancer, but also can be readily applied to other heterogeneous high-throughput data sources. `inteGREAT` is open source and available to download from https://github.com/faryabib/inteGREAT.

## 1 Introduction

Cellular processes are tightly regulated in multiple layers, leading to coordinated function of genes and gene products including transcripts and proteins. Aberrations at each tier of these multilayer regulatory circuits could lead to malignant transformations. It has been shown that combined analysis of data characterizing a variety of biomolecules yields discovery of new insights into tumor biology and facilitates identification of important cancer genes and therapeutic targets [1–4]. These initiatives have increased interest in development of methods for integration of heterogenous data sources [5–7].

The interrogation of information garnered by high-throughput measurements of transcripts or proteins have been used to refine stratification of tumors based on their unique molecular characteristics. Furthermore, analysis of each of these data sources separately has facilitated the discovery of transcript-or protein-based prognostic and diagnostic biomarkers. The salient assumption underlying such comparative studies is that there is a one-to-one relationship between transcript and protein expression [4]. Another implicit assumption is that genome-scale technologies such as next generation sequencing-based transcriptomics and mass spectrometry-based proteomics have comparable sensitivity to capture the activities of these gene products. Yet, examining each aspect of tumor pathobiology alone overlooks potential regulatory mechanisms relating the gene products and the technologies measuring these aspects do not have the same coverage. It is therefore critical to effectively combine information gathered by complementary genome-scale measurements to elucidate common and different molecular features of tumor types.

To address this challenge, methods to integrate heterogeneous data sources such as transcriptomic and proteomic data sets have been proposed. These integration methods range from naive weighted means of transcript and protein abundances [8] to consensus pathways and molecules [9]. Other approaches take advantage of the relationships between gene products to produce a network of associated genes, known as an interactome [10]. *De novo* clustering of interactomes was used to elucidate a subnetwork or pathway containing gene products with functional relatedness [11]. Summarizing the information within each cluster using eigenvectors provides a means to compare clusters [10]. Measuring the network structure between different levels was also proposed as a means for data integration [12]. The disadvantage of grouping gene products is that collapsing these structures into clusters can decrease the sensitivity of biomarker detection. For instance, while a cluster may be classified as clinically significant, an important gene may belong to a different cluster depending on the clustering parameters and algorithm. Grouping gene products also complicates devising gene-centric biomarkers that are the main focus of diagnostic tests. Merging networks to create a summary network was also proposed to address the shortcomings of the clustering-based approaches [13, 14]. These methods of integration were performed on a single phenotype and thus cannot readily identify phenotypic biomarkers differentiating tumor subtypes. We propose that expanding differential expression analysis from the individual level to differential integration can facilitate biomarker discovery.

To this end, we present `inteGREAT`, an algorithm for differential integration. `inteGREAT` generates interactomes for both transcriptomes and proteomes and analyzes their network structures to determine the extent by which a gene product and its related partners are similar across different sources of data while different between cellular phenotypes. Using a framework based on both local and global similarity, `inteGREAT` provides a robust and scalable algorithm that can integrate any number of genomic and functional genomic data sets to identify differentiating tumor biomarkers. `inteGREAT` by design does not cluster gene products at any point in order to retain individual relationships and is thus able to assign confidence of integration to each gene representing its transcript and protein expressions. We assessed the ability of `inteGREAT` to detect perturbations in multiple networks through simulations. Using breast cancer transcriptome and proteome data from The Cancer Genome Atlas (TCGA) [15] and the Clinical Proteomic Tumor Analysis Consortium (CPTAC) [16, 17], we demonstrated the utility of `inteGREAT` to identify subtype-specific biomarkers in breast cancer. `inteGREAT` is a robust, easy to use software package and can be generally applied to any abundance data or pre-made network. `inteGREAT` is open source and available to download from https://github.com/faryabib/inteGREAT.

We further applied `inteGREAT` in a pan-cancer integrative analysis of transcriptome and proteome data sets from TCGA and CPTAC for serous ovarian carcinoma (OV) [2, 18], breast cancers (BRCA) [3, 19], colon (COAD) and rectal (READ) adenocarcinomas [4, 20]. We proposed using a measure of clumpiness on the resulting hierarchy of comparisons that elucidated the promiscuous nature of the luminal and HER2-positive subtypes, while demonstrating the relative isolation of ovarian, colorectal, and to some extent basal subtypes. Our integrative pan-cancer analysis quantitates the importance of each individual gene in stratifying a particular subtype. Among them, we identified a set of clinically important genes that are strongly associated with prognostic outcomes in a given tumor type. Our differential integration of transcript and protein abundance across four tumor types is a showcase of using `inteGREAT` for similar integration analysis in other cancers and diseases.

## 2 Materials and Methods

### 2.1 inteGREAT algorithm overview

`inteGREAT` is an algorithm for integration of disparate high-throughput data sets. This algorithm can perform differential integration for comparative analyses of multiple cellular phenotypes. Differential integration is crucial to stratify two phenotypes and uncover genes leading to molecular differences between tumors. `inteGREAT` achieves differential integration in three stages: network generation, network similarity, and vertex joining (Figure 1). `inteGREAT` first creates an undirected graph of correlations between gene products, called an interactome, for the transcriptome and proteome separately. Each vertex is a gene product and each edge is a correlation between two gene products. `inteGREAT` implements several correlation measures, such as Spearman’s or Pearson’s correlation coefficient. A strong connection requires two gene products to exhibit similar behavior across two cellular states.

**Figure 1:**
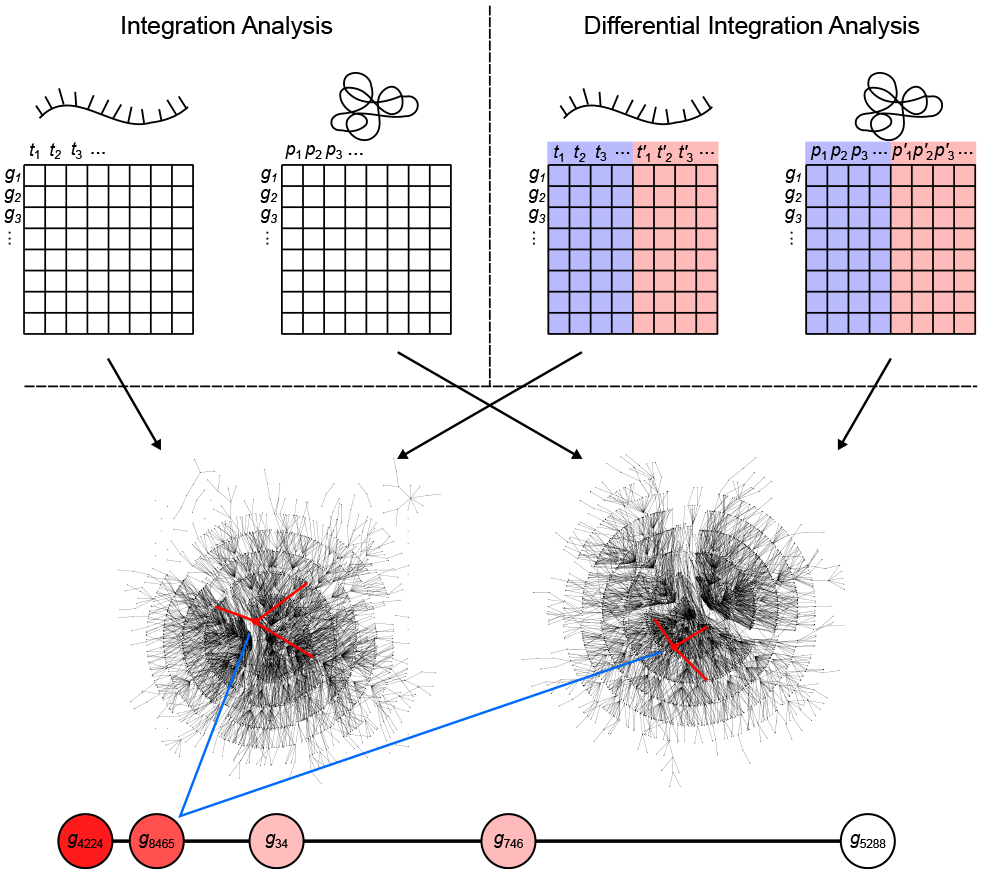
Overview of the `inteGREAT` algorithm. Abundance data for all genes from the transcriptome (left) and proteome (right) provides information to generate a correlation network for each data source. `inteGREAT` measures the local and/or global similarity for each gene and produces a ranked-order list of putative biomarkers with assigned confidences.

To determine vertex-wise network similarity, `inteGREAT` provides two methods. First, `inteGREAT` analyzes the structure of the immediate neighborhood of a gene in each interactome and calculates the cosine similarity between gene products, i.e. interactome vertices. Second, for an expanded measure of topology, `inteGREAT` uses random walk with restart to obtain a stationary distribution centered around a gene product in both interactomes and then determines the concordance in structure using cosine similarity between the two distributions. A random walk with restart provides a more global view of the vertex neighborhood (global similarity), while cosine similarity efficiently compares a vertex’s immediate neighbors across two measurement levels (local similarity). By analyzing the topological structure of each interactome before collapsing protein and transcript measurements into a single gene identifier, we can observe relationships of gene products at each level without loss of information. The value resulting from the network similarity step represents how conserved the neighborhood of interactions for a gene product are between the interactome generated from the transcriptome and proteome assays. The joining step produces a final result as a ranked-order of gene cosine similarities. A differential integration using both tumor types reinterprets this value as conserved behavior across assays but different between the two cellular phenotypes.

### 2.2 Correlation networks

The first step of `inteGREAT` involves generating the correlation matrix where each gene is a vertex and each edge is a correlation. Let *G* be the set of genes. From each gene pair *g*_*i*_, *g*_*j*_ ∊ *G*, we create the correlation network adjacency matrix **A**, where

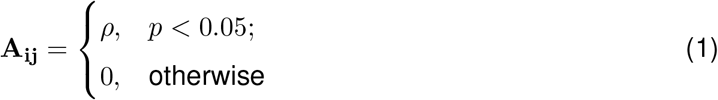

such that *ρ* and *p* are correlation coefficients and *p*-values respectively from any correlation measure between genes *g*_*i*_ and *g*_*j*_, such as Pearson [21] or Spearman [22] correlations. `inteGREAT` can either take as input normalized abundance data and generate these networks or accept pre-made networks.

### 2.3 Vertex comparison

The main function of `inteGREAT` is to relate vertices between networks. Let *U* = {**A**_1_, **A**_2_,…, **A**_1_} be the set of adjacency matrices representing correlation networks of l data sources. We find the integrated value of all genes by assigning each vertex a “vertex similarity score”. We calculate this score by using either cosine similarity [23] with random walk with restart (global similarity) [24] or without (local similarity).

Let the cosine similarity between two vectors of equal length be defined as

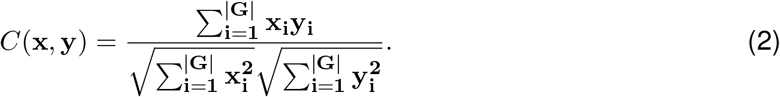

Then for **local** scores, the correspondence vector c between any two adjacency matrices A_x_ and A_y_ is defined as

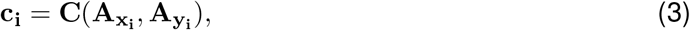

where c_i_ is the correspondence between A_x_ and A_y_ for gene *g*_*i*_.

For **global** scores, c is defined as

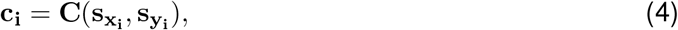

where 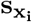 and 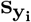 are the stationary distributions for 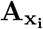 and 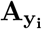 respectively. The stationary distribution of gene *g*_*i*_ in a network with adjacency matrix A_z_ with restart is defined as [24]

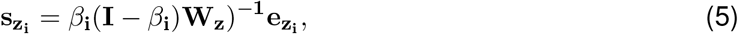

where **I** is the identity matrix, 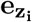 contains 1 at position *i* and 0 elsewhere, and 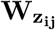 is the probability of traveling from *g*_*i*_ to *g*_*j*_ [24]. **W**_z_ is calculated from **A**_z_, such that 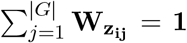, ∀_*i*_, *j* ∊ {1,2,…, |*G*|}, and 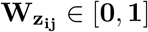.

For *l* > 2, we calculate all c for each pairwise comparison in *U* and use a joining function *f* to combine the values at each *g*_*i*_ into a final value. Function *f* can be the maximum, minimum, arithmetic mean, geometric mean, rank product, etc. So for *l* = 3, we can have 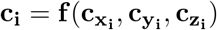, where *f* is the arithmetic mean for instance. In this study, we have two data sources and so there is no need for *f* (outside of simulations), however `inteGREAT` defaults to *f* as the arithmetic mean.

### 2.4 Score confidence

We assign confidence intervals to each score in c using the bias-corrected and accelerated (BCa) bootstrap [25]. As both the global and local similarity scores end with cosine similarity to get the correspondence at c_i_, we can use bootstrapping on the cosine similarity between two vectors at this same step. Let x and y be two vectors of length |*G*|. Then let *α* and *β* be two vectors of length *n* < |*G*|, such that *α*_*i*_ = x_j_ and *β*_*i*_ = y_j_ ∀_*i*_ ∊ {1,2,…, |*G*|} ^ *∀*_*j*_ ∊ {1,2,…, *n*} (so the relationship of indices are maintained from x and y to *α* and *β*). Our bootstrapping function is then *C*(*α*,*β*). We expect the resampled vectors to have a similar direction as the complete vectors. From this analysis we obtain the confidence interval as well as the confidence interval width, the measure we use to assign confidence.

## 3 Results

### 3.1 inteGREAT provides robust measures of inter-network similarity

A key component of differential integration is detecting changes concordantly reflected across multiple gene products associated with a given gene. However, technical or experimental variabilities can lead to noisy high-throughput data sets, potentially resulting in unreliable inference. Missing or inaccurate gene product readouts and variability in the assays can result in interaction networks with missing vertices, misplaced edges, and noisy edge values. As `inteGREAT` detects changes concordant across different data sources using network similarity, we evaluated how unreliable data impacts identifying similarity between networks.

In order to mimic a biological network with hubs, we randomly generating a network using the Barabási-Albert model [26] to represent an interaction network from a single level of data. In the absence of noise, this network represents a single interactome produced by any of the data sources. To simulate interactomes from additional data sources, we generated a new network by permuting 5% of the vertices in the original network. These vertices were the “difference” between data sources and acted as known changes `inteGREAT` attempted to detect. We simulated the scenarios when two or three data sources are available, such as transcriptomics, proteomics, and phosphoproteomics and assessed the performance of `inteGREAT` with global or local similarity was assessed (Figure 2, see Supplementary Materials). To complement the stationary distribution of the random walk with global similarity, we included the result of having simulated random walk transition through the network with restart. As the result of `inteGREAT` is a ranked-order of genes, we measured accuracy by the overall distance of each changed vertex from its expected location at the end of the list (Table S1, see Supplementary Materials).

We first simulated scenarios where noisy measurements result in false relationships between gene products. To this end, we permuted 0% to 50% of edges in the network. This permutation could model differences between the molecular species measured by each high-throughput technology. Regardless of the similarity measure, `inteGREAT` was invariant to the size of the network but became slightly more accurate when the behavior of a cellular system was characterized with three interactomes (Figure 2A), suggesting a potential benefit in investigating a phenotype at multiple data sources − for instance, including not just the transcriptome or the proteome, but the epigenome as well. We also observed similar accuracy for both local and global similarity. `inteGREAT` with local similarity exhibited slightly improved performance when two data sources were considered, suggesting that measuring direct neighbors instead of an interactome global view captured by random walk is more robust when two gene products are associated based on one data source but are independent based on the other vantage point of the system.

We next sought to explore the effect of missing information on our network similarity measures through vertex deletion (Figure 2B). Missing data is common when comparing transcriptome and proteome measurements of the same cellular condition, as the breadth (number of measured proteins) may not encompass all the genes found in the transcriptome analysis. To simulate missing data, we randomly deleted 0% to 50% of the vertices. Among the simulated scenarios, vertex deletion resulted in the worst performance compared to the other sources of noise, suggesting the lack of measurement of a gene product’s neighbor in a data source could not be fully compensated for by observing that neighbor at alternative levels. Integrating data sources from a number of high-throughput technologies with different breadth in measurements significantly limits the accuracy of the integration analyses. Comparison of integration analyses when two or three data sources were available showed that an additional data source enhanced the accuracy of the integration analysis, implying that more comprehensive characterizations of a cellular phenotype using complementary assays could alleviate the detrimental effect of imbalanced breadths of various technologies and missing data.

We also simulated the effect of noisy high-throughput experiments on the accuracy of transcriptomics and proteomics integrative analysis. To simulate this source of network inaccuracy, we injected noise into each edge from a normal distribution with *σ* from 0 to 6 (Figure 2C). This simulation resulted in a striking difference between the `inteGREAT` performance with local and global similarity. The performance of `inteGREAT` with global similarity was minimally impacted by the introduced noise, while `inteGREAT` with local similarity exhibited performance decrease proportional to the noise level. (Figure 2C). Nevertheless, `inteGREAT` performed with greater than 0.8 accuracy, which is significantly higher than the worst-case accuracy of 0.5 resulting from changed vertices uniformly distributed among the ranked-order list. This result suggests that `inteGREAT` can be reliably deployed even in the presence of some degree of inconsistency between networks and is robust to noisy measurements.

**Figure 2:**
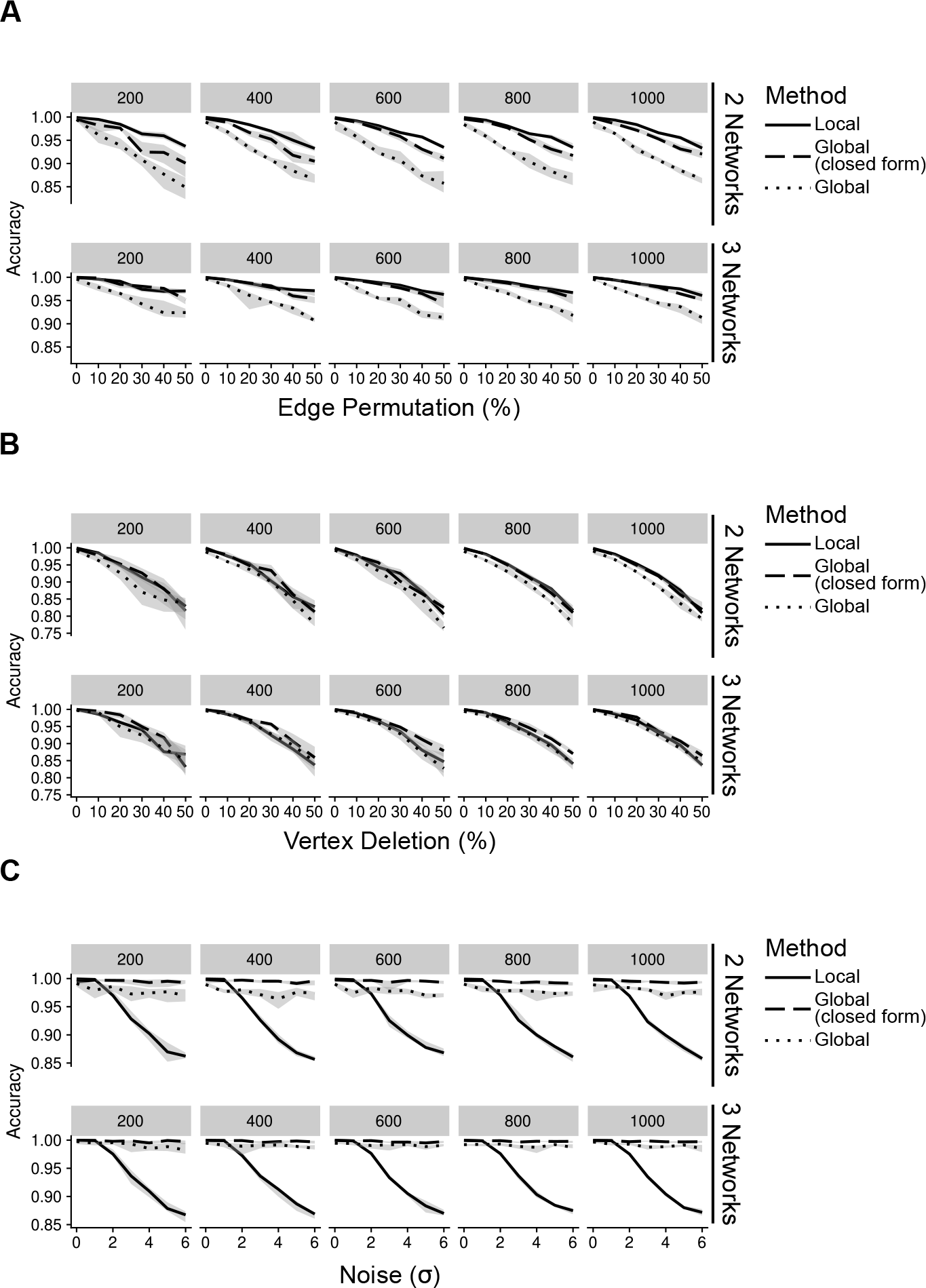
Accuracy of `inteGREAT` for various noise types across different number of data sources (networks) and network sizes (facet label). (A) Simulations with varying percent of the edges permuted. Existing edges are randomly chosen for permuting based on a uniform distribution. (B) Simulations with varying percent of the vertices deleted. Vertices are randomly chosen for deletion using a uniform distribution. (C) Simulations with varying amount of noise. Noise refers to the addition of random values to the network edges drawn from a Gaussian distribution with a standard deviation of the x-axis.

### 3.2 inteGREAT rediscovers canonical biomarkers of breast cancer subtypes

Although integration of synthetic networks demonstrated the robustness of `inteGREAT` in the presence of various sources of noise in the measurements, simulated data are generally limited in recapitulating the complexity of real biological data sets. We next investigated the ability of `inteGREAT` to identify biomarkers associated with a given cellular phenotype from transcriptomic and proteomic data sets.

We conducted differential integrative analyses using TCGA transcriptomic and CPTAC proteomic data sets of basal and luminal breast cancer subtypes [27]. The `inteGREAT` differential integration analysis resulted in a ranked-order list of 13958 gene identifiers (representing respective gene products) from the most to the least conserved between the transcriptome and proteome and differential between the basal and luminal subtypes. We hypothesized that the genes with the most conserved neighborhoods in all data sources and different between the luminal and basal subtypes would be placed at the top of the ranked-order list. To test this hypothesis, we benefited from the curated MSigDB gene set database [28] and performed unbiased gene set enrichment analyses (GSEA) [29] on the ranked-order list outputted by the `inteGREAT` differential integration analysis. The genes identified by `inteGREAT` as highly different between the luminal and basal subtypes while exhibiting concordant transcript and protein neighborhood topologies were significantly enriched with the gene-programs known to differentiate basal and luminal subtypes (Figure 3A, Table S3). These sets included genes that are positively regulated by estrogen receptor ER*α*, genes upregulated after estradiol treatment, and genes reported as differential biomarkers of luminal versus basal subtypes in two independent studies (Figure 3A, Table S3). Conversely, the genes that were ranked low and uncorrelated between the data sources were overrepresented in more general pathways unrelated to the pathobiology of basal, luminal, or breast cancer such as HIV infection or proteasomes (Table S4). This result suggests that `inteGREAT` correctly identified the gene-programs and pathways discriminating between these two breast cancer subtypes from the ones that are irrelevant to this comparative study.

We also assessed the benefit of integrating transcriptomic and proteomic data sources rather than using only one source by comparing the results of integrative and single data source analyses. To this end, we applied `inteGREAT` such that the two interactomes were generated from only the basal or luminal transcriptomic data sets, and used `inteGREAT` to identify the differences between the two transcriptomic networks. We also performed a similar single data source analysis based on the proteomic data sets instead of transcriptomic measurements. These two analyses resulted in two ranked-order lists of genes: one from the transcriptome and the other from the proteome. Compared to the differential integrative analysis, single data source analysis based on the transcriptome or the proteome alone both detected fewer gene sets implicated in the pathobiology of breast cancer and the differences between the basal and luminal subtypes (Figure 3A, Tables S5 and S6). For instance, neither of the single data source analysis were able to identify the genes positively regulated by *ESR1*, a known activated pathway in the luminal subtype. Together, these analyses exhibit the ability of `inteGREAT` differential integration analysis to not only elucidate some of the known gene-programs and pathways associated with the differences between the basal and luminal subtypes, but also underscores the benefit of additional data sources for more accurate integrative analysis.

**Figure 3:**
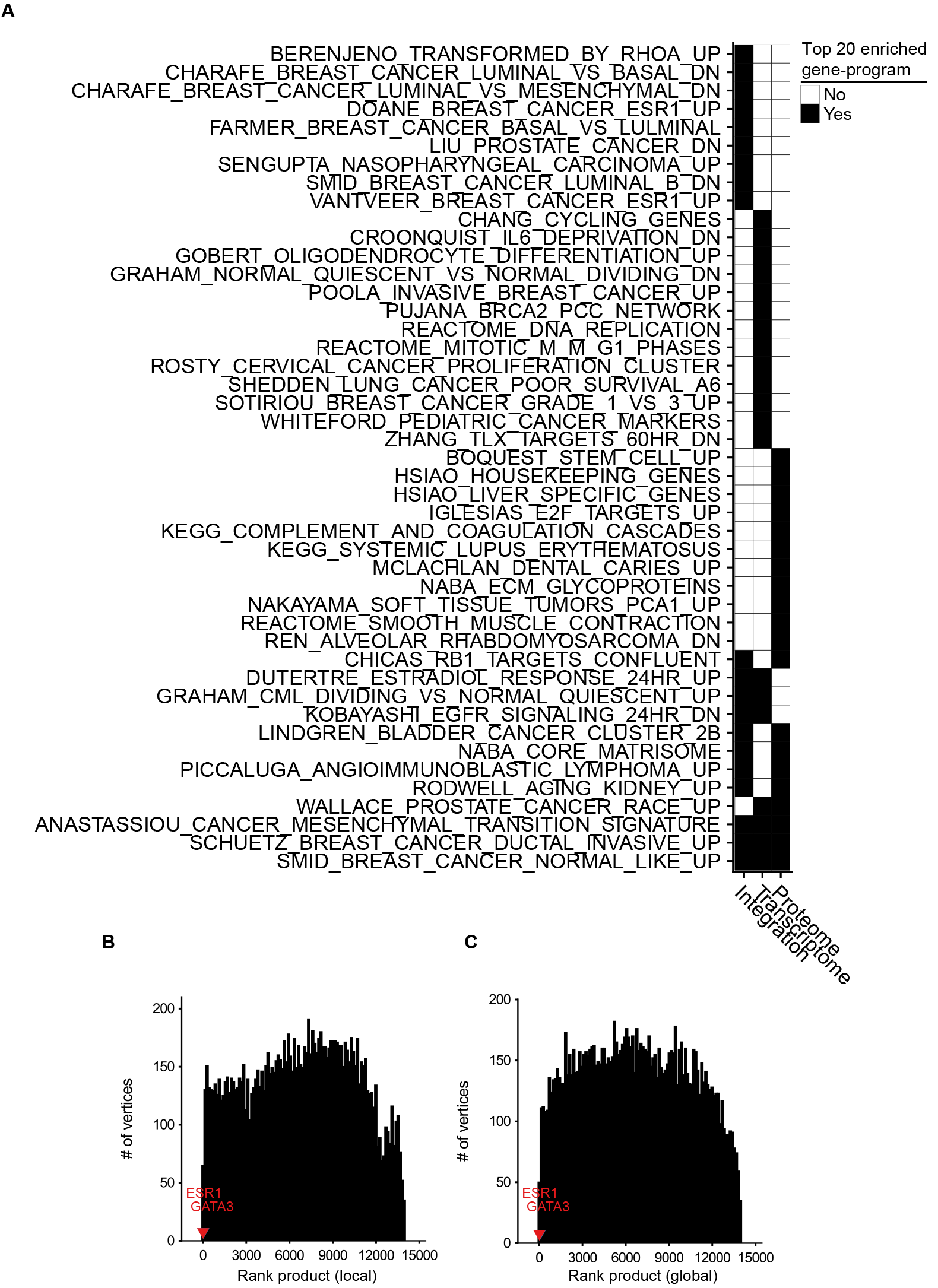
Differential integration of basal vs. luminal breast cancer subtypes identified known gene-programs associated with these tumor subtypes and detected *ESR1* and *GATA3* as their differential biomarkers. (A) Top 20 gene-programs associated with the most differential genes between basal and luminal breast cancer subtypes for three differential analyses: integration of transcriptome and proteome, transcriptome only, and proteome only. Pre-ranked GSEA analysis was performed based on the gene-programs defined by MSigDB C2 curated gene sets and the ranked-order list of differential genes generated by each analysis. Black cells signify a gene set or pathway in the top 20 most significantly enriched pathways for that column. (B) 10 runs of differential integration of basal vs. luminal using local similarity. The final ranked-order list was generated from the joining of each ranked order-lists using the rank product. *ESR1* and *GATA3* are marked with red and blue respectively. (C) Final rank product of 10 runs based on global similarity.

One of the advantages of not collapsing genes to pathways is the direct identification of potential biomarkers of a tumor subtype. *ESR1* and *GATA3* are reported as differential biomarkers of basal and luminal subtypes [27, 30, 31]. In the ranked list of 13958 gene identifiers resulted by the differential integration analysis of basal vs. luminal with local similarity, *ESR1* and *GATA3* were ranked 2nd (CI: 0.377 – 0.407) and 7th (CI: 0.356 – 0.390), respectively. We compared these rankings to integration analysis within each single tumor subtype to assess the benefit of biomarker detection using differential integration. Without differential integration, we observed a marked decrease in the rankings for these known differential biomarkers of breast cancer subtypes. Integration of basal subtype proteomic and transcriptomic data ranked *ESR1* and *GATA3* at 8889 (CI: 0.0100 – 0.0391) and 2754 (CI: 0.0365 – 0.0705), respectively. Integrative analysis of luminal A subtype ranked *ESR1* and *GATA3* at 1688 (CI: 0.0558 – 0.0915) and 9027 (CI: 0.0166 – 0.0483), respectively. While the similar analysis in luminal B, resulted in 133 (CI: 0.181 – 0.220) and 2345 (CI: 0.0607 – 0.0990) ranking of *ESR1* and *GATA3*, respectively. Furthermore, standard differential expression analysis of each data source ranked *ESR1* and *GATA3* at 5 and 128 most differential transcripts respectively. Similar analysis of proteome data set ranked *ESR1* and *GATA3* as 17 and 23 most differential proteins between the luminal and basal subtypes. Together this analysis demonstrates that nominating potential biomarkers by differential integration is in the orders of magnitude more accurate than the integration analysis of each tumor subtype separately and outperforms standard differential fold change analysis, pointing to the benefits of differential analysis.

Potential biases in the collected data sources could adversely impact analyzing the interactome structures separately. For instance, it is common to have more samples from one assay over another. In the TCGA/CPTAC breast cancer data sets, there are 29 more proteome samples compared to transcriptome samples. As a result, there could be some degree of overfitting in a co-expression network construction leading to more accurate inference of the network with a larger sample size. Hence, we postulated that enrichment of proteome samples might have resulted in a bias in our networks. In order to evaluate the robustness of `inteGREAT` to uneven number of samples between the two data sources, we randomly sub-sampled our basal and luminal data sets such that there were an equal number of samples for the transcriptome and the proteome. The similarity and the confidence interval (CI) width was calculated with local (representative run, Figure S1A, C) or global similarity (representative run, Figure S1B,D) for each sub-sampled set. The aggregate ranking of the genes was calculated by combining the results of 10 sub-sampled sets using rank product with 1000 permutations (Figure 3B,C). `inteGREAT` with local similarity ranked *ESR1* and *GATA3* as the 2nd (*p* < 1e-16) and 16th (*p* < 1e-16) most conserved gene products that are at the same time differential between basal vs. luminal subtypes (Figure 3B and Table S5), respectively. A similar analysis using `inteGREAT` with global similarity ranked *ESR1* 3rd (*p* < 1e-16) and *GATA3* 22nd (*p* < 1e-16) (Figure 3C and Table S6). These results corroborate with the expected biomarkers shown to differentiate these two breast cancer subtypes [27, 30, 31], implying the robustness of our framework to biased sample sizes.

### 3.3 inteGREAT relates molecular signatures and tissue-of-origin tumor classification

After establishing the ability of `inteGREAT` to identify differential biomarkers of basal versus luminal breast cancer subtypes, we sought to elucidate relationships between molecular underpinnings of cancer types and their organs of origin. To this end, we expanded our data set to encompass transcriptomic and proteomic data sets for serous ovarian carcinoma (OV) [2, 18], breast cancers (BRCA) [3, 19], colon (COAD), and rectal (READ) adenocarcinomas [4, 20] (Table S2, see Supplementary Materials). We applied `inteGREAT` to each of the malignancies in our data set to assess intra-cancer (e.g. colon transcriptome and proteome) conservation between a gene’s transcript and protein, and evaluated the relationships between cancer types based on the Spearman’s correlation between the `inteGREAT` intra-cancer integration results. The tumors originating from colon and rectal tissues exhibited strong molecular similarities (Figure S2A). Commonalities between colon and rectal samples were previously noted [32]. Our results expanded those findings through the use of only two platforms, one not included in [32]. Breast cancer subtypes classified as luminal A and B based on their transcriptome signature [33] were also significantly correlated (Figure S2A). These observations corroborate with previous work [33], where similarity between mutation, copy number, and DNA methylation of these breast cancer subtypes were reported.

To provide a more refined and quantitative picture of the extent of differences among these tumor classes, we complemented the intra-cancer integration analysis with inter-cancer integration analysis by applying `inteGREAT` to each cancer pair (e.g. colon vs. ovarian transcriptome and proteome data sets). Hierarchical clustering of the intra-cancer and differential integration identified seven distinct clusters (five branches as cut distance 1.54, one of which consists of three branches at cut distance 1.33) and yielded a distinct relationship between their transcript/protein expressions and tissues of origin (Figure 4A). We observed that the BRCA luminal A/B subtypes clustered together. The BRCA basal subtypes were distinct from the luminal subtypes, an observation that was noted earlier by integrative genomics analysis [32]. Ovarian tumors form a distinct cluster which exhibited their differences from BRCA subtypes. Colon and rectal cancers were distinctly identifiable and neighbored the cluster consisting of differences between the ovarian and colorectal tumors. The last two clusters were formed by the differences between the rectal and colon versus breast tumors (Figure 4A). Interestingly, the HER2-positive breast cancer subtype was spread across the dendrogram (Figure 4A). In stark contrast, ovarian cancer was strongly segregated in a single subtree, only appearing elsewhere close to colon and rectal cancers.

In order to clarify the aggregation of ovarian cancer comparisons and the promiscuous placements of HER2-positive breast cancer subtype within the dendrogram, we applied a clumpiness measure [34, 35] to the tree in Figure 4A (see Supplementary Materials). Clumpiness is a measure of aggregation of labels within a hierarchical structure. With this measure, we can quantify the degree of dispersion of a cancer throughout the dendrogram. We observed that ovarian cancer was indeed the least similar to all other cancer types included in our pan-cancer analysis, but shared a stronger relationship with colorectal cancers than the breast cancer subtypes (Figure 4B heatmap). Interestingly, while the colon and rectal cancers were aggregated with themselves, similar to ovarian cancers, the breast subtypes were not aggregated into a single group (Figure 4B heatmap). In fact, HER2-positive and luminal A subtypes had low clumpiness values with themselves, meaning their comparisons were scattered across the entire dendrogram of Figure 4A (Figure 4B heatmap). This finding implies a weak intra-cancer relationship; these tumor types have stronger similarities with other types than their own. Furthermore, by hierarchically clustering these clumpiness values, we observed an overall relationship of cancers (Figure 4B). The dendrogram consists of two distinct groups: the breast cancer subtypes and the colorectal / ovarian subtypes. We found that luminal A and B were the most related and as a sub-group the most different from basal subtype. This observation demonstrates the discrepancy between the luminal and basal cells in the mammary ducts, in line with previous studies [27]. Furthermore, we observed that the HER2-positive subtype was more closely related to luminal than basal subtypes (Figure 4B), possibly because some of the luminal B tumors carry ERBB2 amplifications, while all the tumors classified as basal subtype in our data set are triple negative and lack HER2 expression. We also observed that colon and rectal cancers converged into a colorectal cancer type (Figure 4B), as reported earlier in [32]. This finding reflects the close tissue proximity of the two cancers. Most dissimilar to all other cancer types in our data set was ovarian cancer (Figure 4B), which is known to have a unique signature [36]. Our integrative analysis using `inteGREAT` produced similar relationships between the tumor molecular features and tissue-of-origin as [32], while using measurements from two platforms instead of five (Figure 4B).

**Figure 4:**
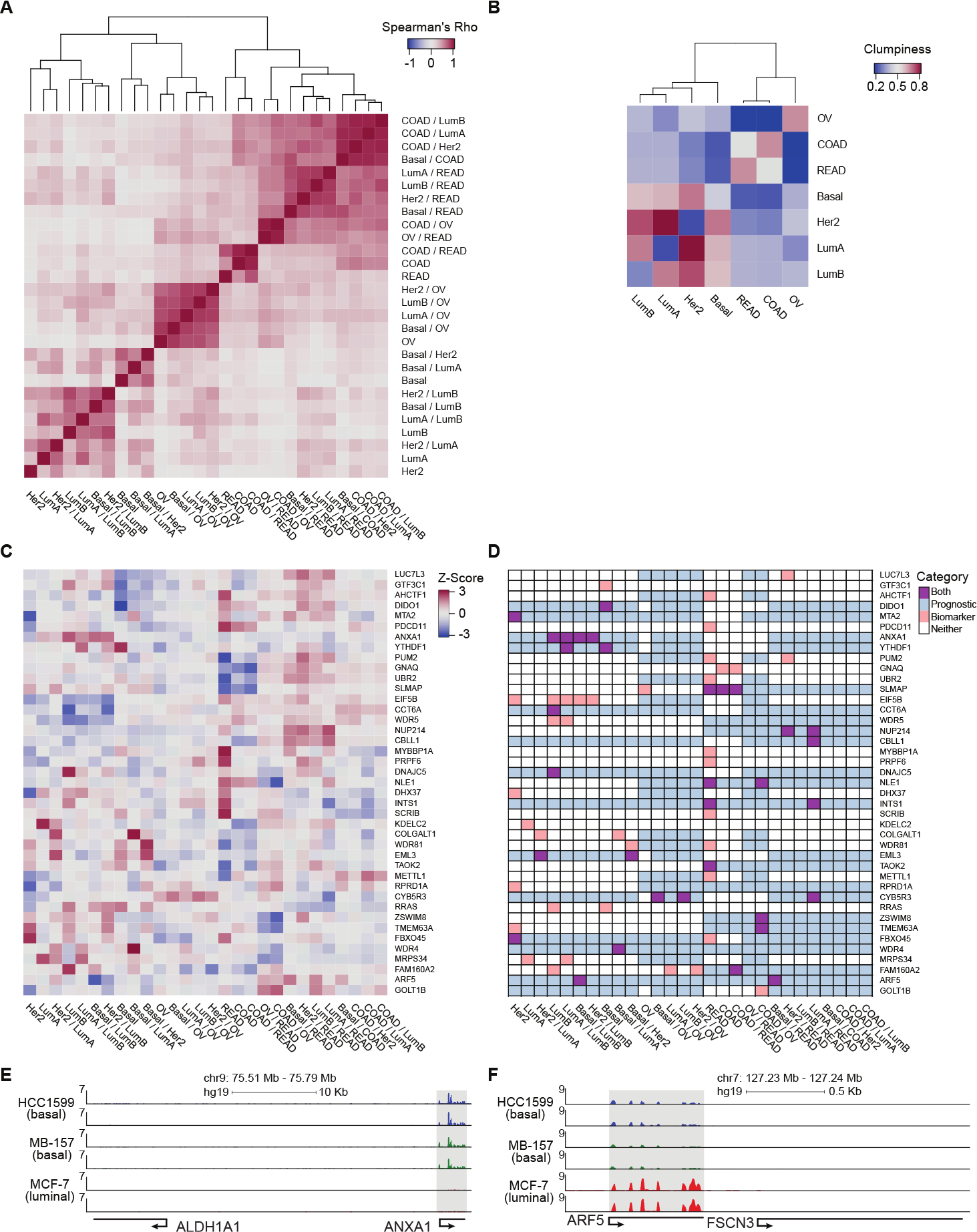
Pan-cancer differential integration. (A) and (B) elucidate relationships between the tumor molecular features and tissue-of-origin. (A) Heatmap of Spearman correlations between local similarities of differential integrations; (B) heatmap of clumpiness values of cancer types from the dendrogram of hierarchically clustered columns of Spearman’s rhos for the differential integrations in (A). (C) Identification of putative diagnostic biomarkers. Heatmap of genes with at least one outlier in a differential integration analysis. Columns underwent *z*-score normalization before outlier removal, rows after removal. Genes with CI widths < 0.04 were removed. (D) Prognostic significance of putative biomarkers from (C) inferred from survival analysis of clinical outcomes reported in the Pathology Atlas database. Each gene was designated as one of four states in each comparative study: an outlier comparison for that gene (orange), significant prognosis of that gene in a tissue in that comparison (blue), both an outlier and prognostic (purple), and neither (white). RNA-seq analysis of *ANXA1* (E) and *ARF5* (F) expression in HC1599 (basal), MB-157 (basal), and MCF-7 (luminal) cell lines.

### 3.4 Pan-cancer differential integration identifies putative prognostic biomarkers

In order to explore the genes acting as possible prognostic biomarkers for each cancer type, we first identified subtype-specific putative biomarker genes. We considered a gene as a putative biomarker if its normalized cosine similarity distribution, generated from the collection of `inteGREAT` intra-and inter-cancer integration analyses, had at least one outlier value, defined as 1.5 times the interquartile range plus or minus the upper and lower quartile, respectively (see Supplementary Materials). An outlier represents a gene that was scored significantly different in one integration analysis compared to the others.

Then we assessed the clinical relevance of these putative biomarker genes. We mined the Pathology Atlas [37] and examined how the expression of our nominated putative biomarker genes correlated with the clinical outcomes as measured by the significance in the differential overall patient survival times for each specific malignancy included in our pan-cancer data set (see Supplementary Materials). The intra-cancer integration analysis identified 93 putative biomarker genes (Figure S2B). The expression level of 38 out of 93 putative biomarkers identified by intra-cancer integration (40.9%) significantly correlated with the differential overall survival rate of cancer patients (Figure S2C). When a similar analysis was performed considering both intra-and intercancer `inteGREAT` analysis, the number of putative biomarkers were reduced to 41, 20 of which (48.8%) exhibited significant correlation with differential overall survival rate (Figure 4D). This observation underscores the importance of differential integration analyses and suggests that finding how much a gene product is conserved within a tumor type but differs from other tumor types can facilitate discovery of clinically relevant biomarkers.

Earlier studies elucidate the significance of a number of biomarkers nominated by `inteGREAT` to the pathobiology of their corresponding disease. For example, *CBLL1*, or *HAKAI*, is a protooncogene implicated in colorectal cancers [38]. *MYBBP1A* is known to bind and activate p53 and is involved in colorectal cancers [39–42]. Our predicted ovarian specific biomarker *CYB5R3* is reported to be involved in ovarian cancer [43]. Together, our analysis suggests that integration of differential transcriptome and protein data sets improves the specificity of biomarker identification.

Using `inteGREAT`, we also identified *ANXA1* and *ARF5* to be putative biomarkers for basal and luminal breast cancer subtypes with potential prognostic significance. High expression of *ANXA1* promotes metastasis of basal-like tumors and associates with poor prognosis in this breast cancer subtype [44, 45]. To verify *ANXA1* as a biomarker for basal vs. luminal subtypes, we measured mRNA in expression three cell lines: HCC1599 (basal), MB-157 (basal), and MCF-7 (luminal). As predicted by the `inteGREAT` pan-cancer analysis, *ANXA1* exhibited significantly higher expression in the two basal cell lines HCC1599 and MB-157 (Figure 4E). Furthermore, it has been previously shown that *ANXA1* has lower expression in luminal than basal tumor types [44], confirming its identification as a biomarker by `inteGREAT` [44]. Conversely, our RNA-seq analysis of breast cancer cell lines confirmed that *ARF5* is highly expressed in MCF-7 luminal cells, but not expressed in basal cell lines (Figure 4F). These data, together with our pan-cancer analysis, propose *ARF5* as a possible biomarker of luminal breast cancer subtype which has a tumor subtype-specific gene-program in transcript and protein with potential prognostic significance.

## 4 Discussion

High-throughput assays have enabled global profiling of different aspects of tumor characteristics, from the transcriptome to the proteome. A significant step toward more effective cancer treatment is to leverage diverse genome-scale data sources to complement investigation of tumor characteristics. It has been shown that the holistic and integrated views of cancer facilitate discovery of molecular-based diagnostic and prognostic biomarkers and guide precise clinical management and therapeutic decision-making. Yet, while recent algorithms attempt to integrate data sources for individual tumor types, there are still unmet needs for analytic approaches to enable differential integration analyses and facilitate the discovery of tumor-specific biomarkers from an integrative view of tumor biology. Here, we have presented `inteGREAT`, an algorithm to integrate transcript and protein abundance data and detect differential biomarkers between multiple cancer subtypes.

We have shown the robustness of `inteGREAT` using simulations controlling for multiple sources of biological noise. In addition, we demonstrated the accuracy and utility of `inteGREAT` to infer differences and similarities of four tumor types. `inteGREAT` confidently identified previously published diagnostic biomarkers of basal and luminal breast cancer subtypes from their respective transcriptomic and proteomic data. Using a measure of clumpiness for summarizing hierarchical trees, `inteGREAT` performed differential integration for seven different cancer subtypes and detected convergence and divergence of tumors from various tissues-of-origin according to their transcriptomic and proteomic characteristics. Furthermore, `inteGREAT` identified putative biomarkers for each subtype with potential prognostic significance.

Using multiple analyses, we demonstrated that integration of transcriptome and protein interactomes enhances reliability of biomarker discovery rather than using only each of these measurements alone. We propose that measuring biological systems from more than one perspective diminishes the effect of missing data and noisy assays, while simultaneously elucidating new relationships between disparate data sources that cannot be captured in a single assay.

`inteGREAT` is a generic algorithm for measuring inter-network similarity and is able to report differential information. While in this study we only used `inteGREAT` for biomarker detection from transcriptome and proteome data in different cancer subtypes, our flexible implementation of `inteGREAT` enables new analysis of networks from a variety of biological sources, including the epigenome, CNVs, and mutation data. This algorithm is a powerful tool to further cancer biomarker discovery to aid in therapeutics advancements.

## 5 Conflict of Interest Statement

The authors declare no competing or conflict of interests.

## 6 Author Contributions

RBF and GWS designed experiments. GWS and RBF designed the algorithm. GWS implemented the software, collected and organized data. JP and YZ performed and analyzed sequencing experiments. RBF and GWS wrote the manuscript with comments from all authors. RBF conceived the project, administrated the experiments and analyses, and provided expert advice.

## 7 Funding

This work was supported in part by T32-CA009140 to GWS, LLS-5456-17 to JP and Abramson Family Cancer Research Institute Investigator Award, Abramson Cancer Center Cooper Award, Institute for Translational Medicine and Therapeutics program for Transdisciplinary Awards Program in Translational Medicine and Therapeutics to RBF.

## 8 Acknowledgments

We are grateful to Drs. Kojo Elenitoba-Johnson, Warren S. Pear, and Golnaz Vahedi, Ethan Mack for indispensable advice and critically reading the manuscript. We thank members of the Vahedi, and Pear labs particularly Georgios Georgakilas, Stanley Cai, and Ethan Mack.

## 9 Data Availability Statement

The data sets analyzed for this study can be found in The Cancer Genome Atlas (TCGA) https://portai.gdc.cancer.gov/, the Clinical Proteomic Tumor Analysis Consortium (CPTAC) https://cptac-data-portal.georgetown.edu/cptacPublic/, and the Pathology Atlas https://www.proteinatlas.org/pathology.

